# Membrane fusion reactions limited by defective SNARE zippering or stiff lipid fatty acyl composition have distinct requirements for Sec17, Sec18, and adenine nucleotide

**DOI:** 10.1101/2024.11.15.623832

**Authors:** Karina Lopes, Amy Orr, William Wickner

## Abstract

Intracellular membrane fusion is catalyzed by SNAREs, Rab GTPases, SM proteins, tethers, Sec18/NSF and Sec17/SNAP. Membrane fusion has been reconstituted with purified vacuolar proteins and lipids to address 3 salient questions: whether ATP hydrolysis by Sec18 affects its promotion of fusion, whether fusion promotion by Sec17 and Sec18 is only seen with mutant SNAREs or can also be seen with wild-type SNAREs, and whether Sec17 and Sec18 only promote fusion when they work together or whether they can each work separately. Fusion is driven by two engines, completion of SNARE zippering (which does not need Sec17/Sec18) and Sec17/Sec18-mediated fusion (needing SNAREs but not the energy from their complete zippering). Sec17 is required to rescue fusion that is blocked by incomplete zippering, though optimal rescue also needs the ATPase Sec18. ATP is an essential Sec18 ligand, but at limiting Sec17 levels Sec18 ATP hydrolysis also drives release of Sec17 without concomitant *trans*-SNARE complex disassembly. At higher (physiological) Sec17 levels, or without ATP hydrolysis, fusion prevails over Sec17 release. Stiff 16:0, 18:1 fatty acyl chain lipids provide an alternative route to suppressing fusion, with entirely wild-type SNAREs and without impediment to zippering. In this case, Sec17 and Sec18 restore comparable fusion with either ATP or a nonhydrolyzable analog. Fusion blocked by impaired zippering can be restored by concentrated Sec17 alone (but not by Sec18), while fusion inhibited by stiff fatty acyl chains is partially restored by Sec18 alone (but not by Sec17). With distinct fusion impediments, Sec18 and Sec17 have both shared roles and independent roles in promoting fusion.

**Significance:** Sec17/SNAP and Sec18/NSF catalyze *cis*-SNARE complex disassembly through ATP hydrolysis, but also drive fusion itself without ATP hydrolysis. Fusion can be inhibited by either “stiff” fatty acyl chains while allowing full zippering of entirely wild-type SNAREs or by incomplete SNARE zippering. In either case, Sec17 and Sec18 restore full fusion, but with clear differences. With defective zippering, high levels of Sec17 can restore fusion without Sec18. With stiff fatty acyl chains, Sec18 alone can restore fusion but Sec17 alone will not. In both cases, Sec17 and Sec18 are more efficient together. SNARE-dependent membrane fusion thus relies on two energy sources, complete SNARE zippering and interactions with Sec17 and Sec18.

---

Endocytic and exocytic vesicular trafficking are fundamental for cell growth, protein secretion, neurotransmission, and microbial invasion. Fusion between vesicles and their target organelle membrane needs Rab-family GTPases (Hutagalung and Novick, 2011), Rab-effector tethers (Baker and Hughson, 2015), SNARE proteins (Jahn and Scheller, 2006), SM (Sec1/Munc18)-family catalysts of SNARE assembly (Baker et al., 2015; Orr et al., 2017; Jiao et al., 2018; Stepien et al., 2022), and the SNARE-associated chaperones Sec17/SNAP and the AAA family ATPase Sec18/NSF (Söllner et al., 1993). Organelle identity relies on selective membrane targeting and fusion, itself reliant on the combinatorial match of Rab, SM, and SNARE proteins (Jun and Wickner, 2019). SNAREs have cannonical SNARE domains of approximately 60 amino acyl residues with heptad-repeat apolar residues (Fasshauer et al., 1998; Sutton et al., 1998). SNARE domains are inherently random coil, but become alpha-helical as they zipper in an N-to C direction into a 4-helical coiled coils structure (Sorenson et al., 2006). SNARE domains are flanked by globular N-domains, C-terminal juxtamembrane regions, and (frequently) transmembrane anchors. Each SNARE domain has a central conserved polar arginyl (R) or glutaminyl (Q) residue instead of an apolar residue. SNAREs are in conserved R, Qa, Qb, and Qc families (Fasshauer et al., 1998), and assemble into RQaQbQc complexes, in *cis* (if anchored to the same membrane) or *trans* (if anchored to separate apposed membranes). SNARE assembly is controlled by selective tethering by Rabs and tethers, then catalyzed by SM family protein associations with the R– and Qa-SNAREs on tethered membranes (Baker et al., 2015). SNARE zippering is driven by burying the heptad repeat apolar residues into the interior of the 4-helical coiled coils structure away from water (Tanford, 1980). Complete zippering draws the membranes even closer, and is opposed by the spring-like energy from the resulting membrane bending. Other chaperones bind SNAREs besides SM proteins, such as Sec18/NSF and Sec17/SNAP (Söllner et al., 1993). SNAREs assemble with NSF and SNAP into a 20s complex of known structure (Zhao et al., 2015), and this structure can be envisioned between two membranes in *trans* (Figure 1). The four SNARE domains form a central 4-helical coiled coil, anchored in the two apposed membranes (Figure 1, red, green, blue, tan). A layer of several Sec17/SNAP molecules (yellow) surrounds the SNAREs, each with two leucines (LL) at its C-terminal end to form the site of Sec18/NSF binding near the N-terminal ends of the SNARE domains. The N-domain of Sec17 is near the membrane surface, and a small apolar loop in this Sec17 N-domain is poised to directly engage the bilayer. As envisioned in cross-section, (Figure 1, bottom), basic residues in the center of Sec17 bind ionically to acidic residues in the SNARE domains. The two edges of Sec17 which face other Sec17s surrounding the SNAREs are acidic on one side and basic on the other. The asssembled Sec17s which fully surround the SNAREs are positioned for pairwise ionic interactions along each Sec17 edge (Figure 1, bottom, circled charges). Each of these associations confers stability on the 20s structure.

**Figure 1.**
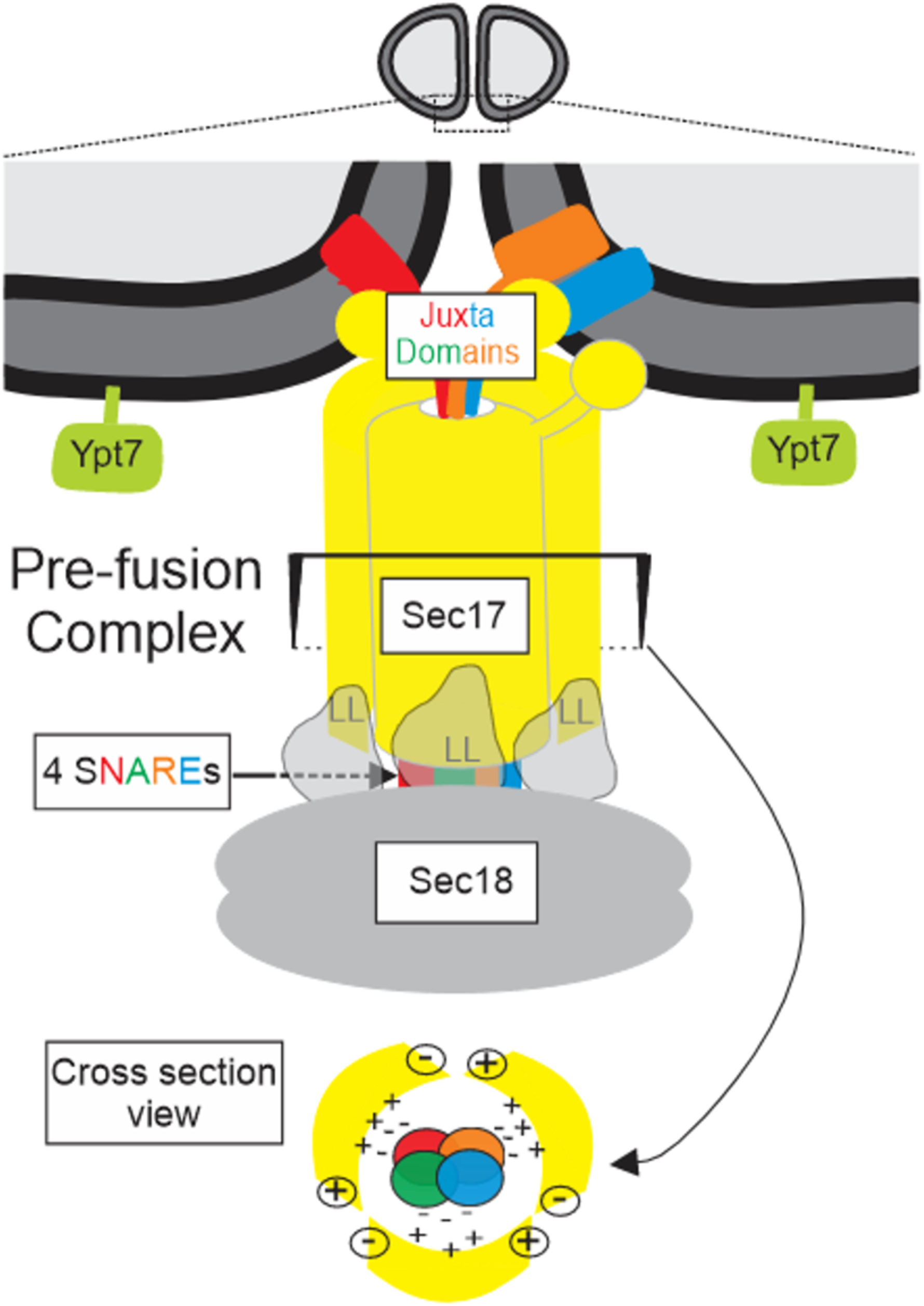
Working model of the trans complex of SNAREs, Sec17 and Sec18. See text for details.

We study membrane fusion with components from yeast vacuoles (lysosomes). Vacuoles undergo fission to form smaller vacuoles and these fuse to form the large, visible organelle often seen at steady state. Wada and colleagues screened for mutants with multiple small vacuoles, identifying 9 nonessential *VAC* genes which encode proteins that exclusively catalyze vacuole fusion (Wada et al., 1992). Other proteins of vacuole fusion which also catalyze fusion along the exocytic pathway are essential for cell viability and therefore escaped the *VAC* screen. The VAC screen identified two of the vacuolar SNAREs, the vacuolar Rab Ypt7 (simply termed Rab hereafter; Haas et al., 1995), and 6 large proteins which are subunits of a protein termed HOPS (**<u>ho</u>**motypic fusion and vacuole **<u>p</u>**rotein **<u>s</u>**orting; Seals et al., 2000). The vacuolar R-SNARE Nyv1, Qa-SNARE Vam3, Qb-SNARE Vti1, and Qc-SNARE Vam7 (termed simply R, Qa, Qb, and Qc hereafter) are on the membrane of each vacuole, but form *trans*-SNARE complexes between R on one membrane and Qa, Qb, and Qc on its apposed fusion partner (Fukuda et al., 2000). Two HOPS subunits which are at the ends of the extended HOPS structure (Shvarev et al., 2022), Vps39 and Vps41, bind the Rab on each fusion partner to effect tethering (Brett et al., 2008; Brocker et al., 2012). The HOPS subunit Vps33 is the SM protein of this organelle and has binding grooves for the N-terminal regions of the R and Qa SNAREs (Baker et al., 2015). When HOPS is bound on one membrane to the Rab, R, and phosphatidylinositol-3-phosphate, it is activated to bind the Q-SNAREs in *trans* and catalyze the assembly of their N-terminal regions with R (Song et al., 2021; Torng et al., 2020). Zippering proceeds N to C, opposed by the energy needed for membrane distortion. Sec17 is enriched on membranes by its assembly with Sec18 into a 3 Sec17:Sec18 complex which binds lipid membranes by product of the lipid affinities of the apolar loop of each Sec17 N-domain (Orr and Wickner, 2022). The binding of this complex to SNAREs may displace HOPS (Collins et al., 2005; Schwartz et al., 2017). The bound Sec17 supports fusion in two ways, associating with SNAREs to promote their zippering (Ma et al., 2016; Song et al., 2021) and providing independent energy for lipid rearrangement through SNARE-proximal insertion of its N-domain apolar loop into the bilayer (Song et al., 2021). Our working model of the pre-fusion complex is shown in Figure 1, based on the pioneering 20s structure of Zhao et al. (2015). The R-SNARE (red) and the Qa (tan), Qb (blue), and Qc (green) SNAREs, anchored in apposed membranes, are surrounded by a shell of Sec17 (yellow). The apolar loop of the N-domain of each Sec17 is inserted into membranes, while two C-terminal leucyl residues bind hexameric Sec18. In cross section (Figure 1, bottom), acidic SNARE domain residues are positioned to form ionic bonds with basic residues of Sec17. Each Sec17 has a basic and an acidic “edge” which are adjacent in the assembled structure (Zhao et al., 2015).

The only known function of ATP hydrolysis in membrane fusion is Sec18-driven disassembly of post-fusion *cis*-SNARE complexes (Söllner et al., 1993; Mayer et al., 1996; Ungermann et al., 1998; Cipriano et al., 2013). Sec18 can disassemble *trans*-SNARE complexes as well, though this is inhibited by HOPS (Xu et al., 2010). Sec18 also promotes Sec17/Sec18-mediated fusion without needing ATP hydrolysis (Zick et al., 2015; Song et al., 2017; Song et al., 2021). We now address three questions: whether ATP hydrolysis by Sec18 can affect its role in promoting fusion, whether fusion promotion by Sec17 and Sec18 is only seen with mutant SNAREs or is seen with wild-type SNAREs as well, and whether Sec17 and Sec18 can only promote fusion when they work together or whether they can each work separately. We find that ATP hydrolysis inhibits Sec17/Sec18-mediated fusion at limiting Sec17 levels but not at physiologically higher Sec17 levels. With limiting Sec17, either 1 mM ADP or ATPγS support Sec17/Sec18-mediated fusion much better than ATP. Surprisingly, fusion inhibition from ATP hyrolysis is not from SNARE complex disassembly, measured by the association of R and Qa in *trans*, but is accompanied by Sec18 ATP hydrolysis-driven release of Sec17 from *trans*-SNARE complexes. Incompletely zippered SNAREs may have fewer sites of SNARE:Sec17 association, allowing facile Sec17 displacement when Sec18 hydrolyzes ATP but not transmitting sufficient force to the SNAREs for their disassembly. Higher levels of Sec17, complete zippering, or limiting ATP levels bypass this sensitivity to Sec18 ATP hydrolysis. Even with entirely wild-type SNAREs, fusion can also be limited by a stiff membrane fatty acyl phase, rich in palmitoyl and oleoyl fatty acyl chains, and this fusion is also restored by Sec17 and Sec18. This restoration is supported equally by ATP or its nonhydrolyzable analogs. Sec17 alone (but not Sec18) can partially restore fusion that was impaired by defective zippering, while Sec18 alone (but not Sec17) stimulates fusion which is limited by stiff fatty acyl chains. In either case, and even with wild-type SNAREs and fluid lipid membranes, optimal fusion requires both Sec17 and Sec18.

## Results

Proteoliposomes were prepared with fluid vacuolar mixed lipids including di-18:2 PC, PE, PA, and PS as well as ergosterol, PI, DAG, and PI3P. Proteoliposomes with either Rab, R, and entrapped biotin-phycoerythrin or with Rab, Qa, Qb, and entrapped Cy5-streptavidin were prepared by detergent dialysis from mixed micellar solutions. For each, the Rab was at a 1:8000 molar ratio to lipids and the SNAREs were at a physiological (Zick et al., 2014) 1:32,000 molar ratio to lipids. In mixtures of these proteoliposomes, the two entrapped fluorphores are kept apart by at least the thickness of two lipid bilayers, too far for fluorescent resonance energy transfer (FRET). Fusion incubations contained HOPS, the Qc SNARE (which is not membrane anchored), and (where indicated) Sec17, Sec18, and adenine nucleotide. Fusion mixes the proteoliposomal lumenal contents. Upon mixing, the binding of biotin to streptavidin brings their attached Cy5 and phycoerythrin fluorophores into intimate proximity, creating a strong FRET signal which is monitored as a measure of fusion (Zucchi and Zick, 2011). All incubations had a large molar excess of nonfluorescent streptavidin to complex with any released biotin-phycoerythrin, preventing FRET due to lysis. With this assay, we measured the effect of hydrolyzable and nonhydrolyzable adenine nucleotides on fusion driven by two means, SNARE zippering (without energetic contribution from Sec18 and the Sec17 apolar loops) and Sec18/Sec17 (without energy from zippering).

With HOPS and full-length Qc, optimal fusion needs both Sec17 and Sec18 but does not require ATP hydrolysis, as it is supported by ATPγS (Figure 2A, filled squares vs open circles; Fig 2B, initial rates, bar 1 vs 4). When apolar residues of the Sec17 N-domain apolar loop are changed to polar serine (Sec17F22SM23S, termed Sec17FSMS), stimulation is largely lost (Figure 2B, bar 2 vs 4 and 6). The apolarity of the Sec17 N-domain loop is needed both for the interdependent delivery of Sec17 and Sec18 to membranes (Orr and Wickner, 2022) and for fusion stimulation once they are at the membrane (Song et al., 2024). Sec17 can promote SNARE zippering (Ma et al., 2016; Song et al., 2021) and induce fusion without energy from completion of zippering (Song et al., 2021).

**Figure 2.**
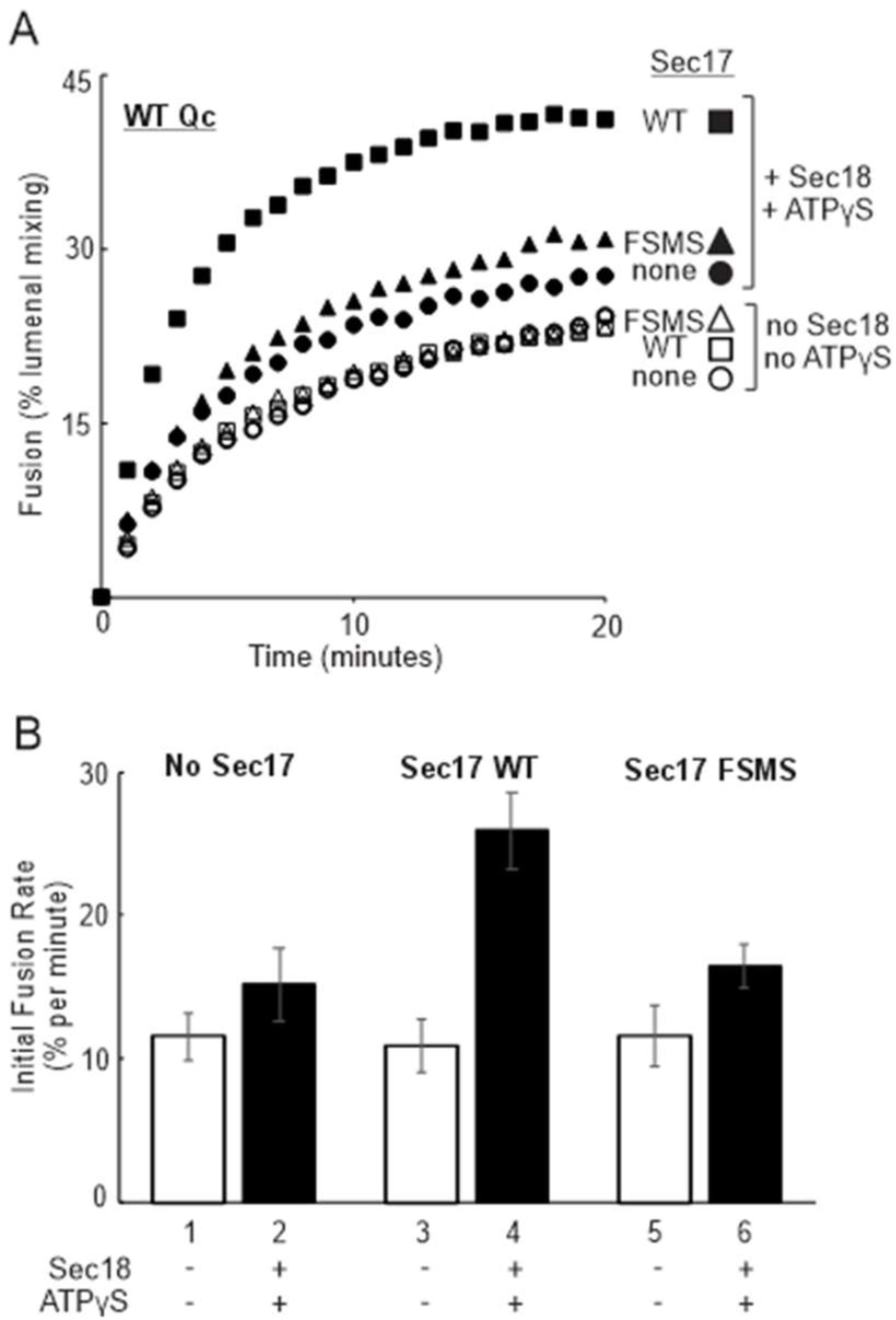
Stimulation by Sec17, Sec17FSMS, and Sec18 of the fusion of proteoliposomes bearing wild-type SNAREs. As described in Methods, fusion incubabations had 50 nM HOPS, 400 nM wild-type Qc, and where indicated had 100 nM Sec17 or Sec17FSMS and/or 300 nM Sec18 with 1 mM MgATPγS.

### Adenine nucleotides and fusion without energy from zippering

Qc3Δ, the deletion of the C-terminal third of the Qc-SNARE domain, blocks zippering-driven fusion but allows fusion driven by Sec17 and Sec18 (Song et al., 2021). High levels of Sec17 (1 μM) are needed to support this fusion without Sec18 (Figure 3A, open diamonds and triangles and 3E, black bars 13, 17). Sec18 and ATP stimulate fusion at 0.3 μM Sec17 or 1 μM Sec17 (Figure 3B; Figure 3E, blue bars 14 and 18). ADP (Figure 3C; Figure 3E, purple bars 4, 7, 11, 15) or ATPγS (Figure 3D; Figure 3E, green bars 3, 5, 8, 12, 16) support Sec17/Sec18-mediated fusion at substantially lower Sec17 concentrations.

**Figure 3.**
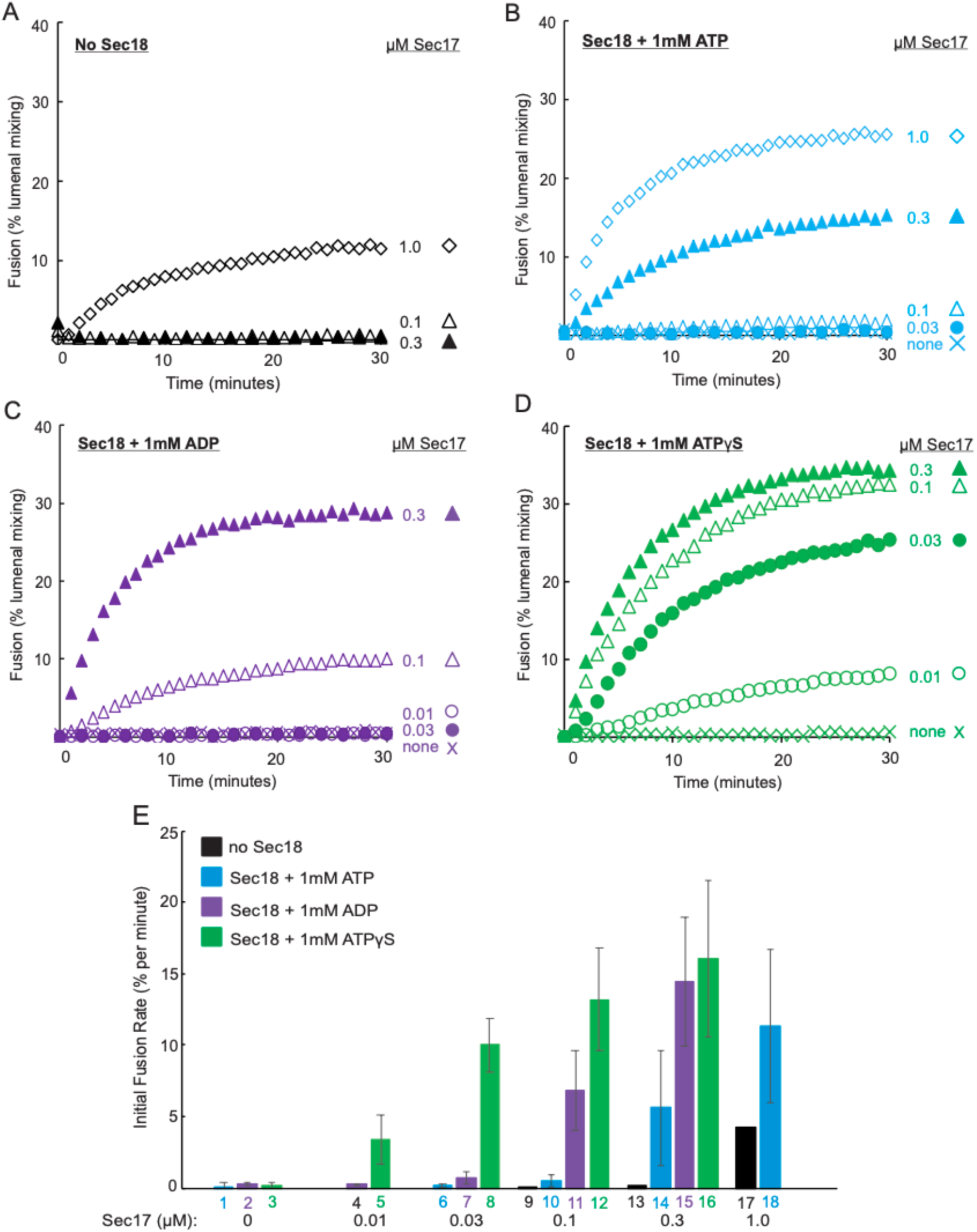
Effects of Sec18 and adenine nucleotide on Sec17/Sec18-mediated fusion. Fusion as in Methods and Figure 1, but with Qc3Δ instead of Qc. Fusion indubations had (A) 1 mM MgATP but no Sec18, (B) 1 mM MgATP and 300 nM Sec18, (C) 1 mM MgADP and 300 nM Sec18, or (D) 1 mM MgATPγS and 300 nM Sec18. In each case, fusion was assayed without Sec17 or with 0.03, 0.1, 0.3 μM, or 1.0 μM Sec17. (E) Initial fusion rates from minutes 1 to 5 of incubation were determined and are shown as means with standard deviations from three repeat assay sets.

What is the basis of the poor fusion with ATP, the physiological ligand to Sec18, with limiting Sec17? There is at least as much interdependent binding of Sec17 and Sec18 to lipid membranes with ATP (Figure 4, A and B, lanes 4, 6) as with ATPγS (lanes 3, 5), suggesting that the interdependent binding of Sec17 and Sec18 to membrane lipid (Orr and Wickner, 2022) is not the basis of Sec17/Sec18-mediated fusion sensitivity to ATP (Figure 3).

**Figure 4.**
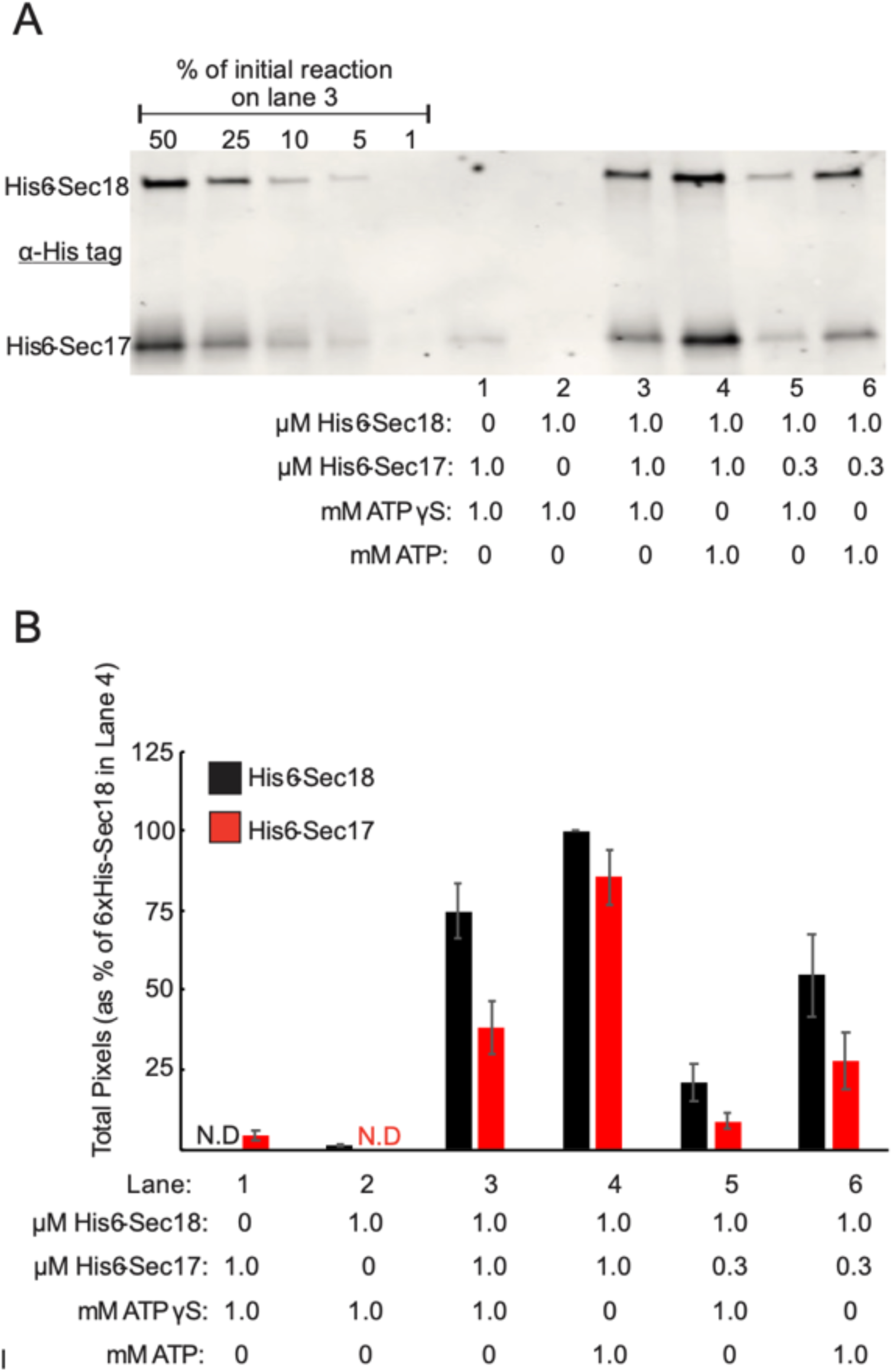
The interdependent association of Sec17 and Sec18 with liposomes is seen with either ATP or ATPγS. (A) Lipid binding assays were conducted according to Orr and Wickner (2022) with 1 mM Mg:ATPγS or Mg:ATP, 1 μM His_6_-Sec18 and 0.3 or 1 μM His_6_-Sec17. Samples were floated and analyzed by Western blot using THE^TM^-His antibody (GenScript USA, Inc., Piscataway, NJ) for the His_6_-tag on the N-terminal region of each protein. The first 5 lanes represented the percentage of input before floatation of sample shown in lane 3. (B) Experiments were repeated three times. Immunoblots were scanned and quantified using UN-SCAN-IT gel 6.3 software. Value of pixels were normalized to the pixels of the His_6_-Sec18 band (black, lane 4).

Sec18 or its mammalian homolog NSF bind ATP but require magnesium for ATP hydrolysis (Zhao et al., 2015). To determine whether ATP hydrolysis accounts for the striking difference between ATP and ATPγS in Sec17/Sec18-mediated fusion at limiting Sec17 (Figure 3), we analyzed fusion with magnesium to allow ATP hydrolysis or in the presence of the magnesium chelator EDTA to block hydrolysis. Sec17/Sec18-mediated fusion with Sec18 and ATP (Figure 5A, open symbols) is stimulated by EDTA (filled symbols) at each concentration of Sec17. This EDTA stimulation is specific to fusion with Sec18 and ATP. With ATPγS, EDTA can even inhibit fusion (Figure 5B, filled symbols with EDTA vs open symbols without EDTA). The average and standard deviations of the initial fusion rates for triplicate experiments are shown in Figure 5C. In the presence of EDTA, Sec18-mediated fusion is comparable with 1 mM ATP or ATPγS (Figure 5C, bars 8 vs 10 and 12 vs 14) but without EDTA, there is far less fusion with ATP (Figure 5C, bars 7 vs 9 and 11 vs 13). With limiting Sec17 levels such as 0.1 μM, fusion is inhibited by Sec18 ATP hydrolysis per se.

**Figure 5.**
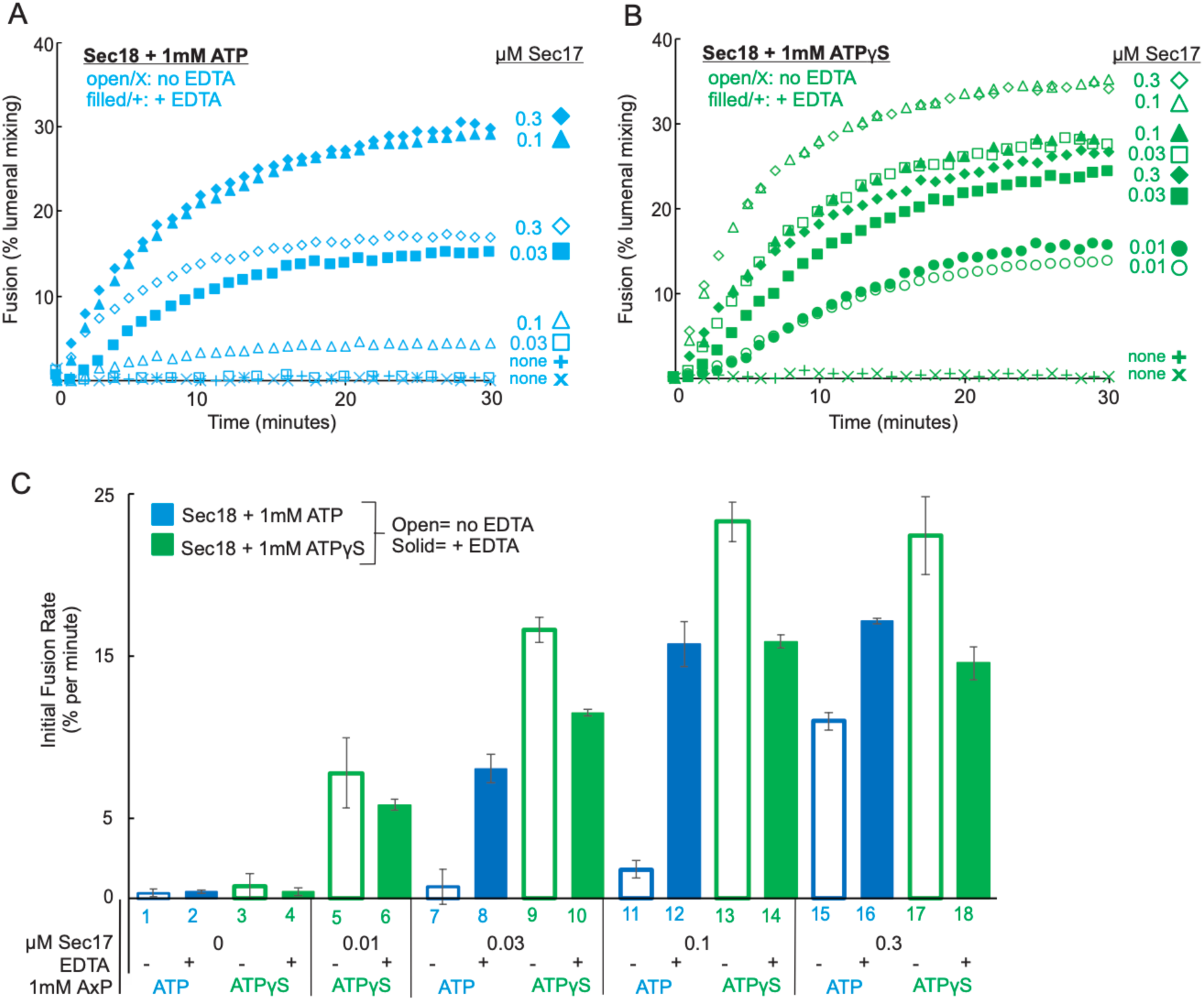
At limiting Sec17, EDTA rescues Sec17/Sec18-mediated fusion with ATP. Fusion incubations had 50 nM HOPS, 40 nM Qc3Δ, the indicated concentrations of Sec17, 1 mM MgATP or ATPγS, 300 nM Sec18, and either no EDTA (open symbols) or 4.5 mM EDTA (filled symbols). Averages and standard deviations of the initial fusion rates are shown for triplicate assays; open bars, no EDTA, filled bars, with EDTA.

### Dual roles of ATP

To determine the conditions where ATP promotes or inhibits Sec17/Sec18-mediated fusion, assays were performed at varying nucleotide concentrations. With defective zippering, fusion needs adenine nucleotide (Figure 6A, x is no adenine nucleotide) and is supported by even 0.025 mM ATP (black circles) or ATPγS (blue circles). Fusion with ATP is optimal at 0.1mM (open black squares), but higher concentrations of ATP progressively inhbit fusion, while increasing ATPγS has no such inhibitory effect (Figure 6A). Throughout this range of ATP or ATPγS concentrations, there is more fusion with ATPγS (blue symbols). With 0.025 mM ATP or ATPγS, fusion is inhibited by EDTA (Figure 6A, open black and blue circles vs Figure 6B, filled black and blue circles, all from the same experiment). Figure 6C shows the initial fusion rates. EDTA inhibits fusion with low levels of either adenine nucleotide (lanes 2 and 3 vs 10 and 11). Nonetheless, when Sec18 ATP hydrolysis is blocked by EDTA, fusion is as well supported by ATP as by ATPγS (Figure 6B, black vs blue symbols, and Figure 6C, bars 10-17), acting simply as a ligand to Sec18 to promote Sec17/Sec18-mediated fusion. ATP hydrolysis by Sec18 inhibits fusion (Figure 6A and Figure 6C, bars 4, 6, and 8). This fusion inhibition by 1 mM ATP is bypassed by EDTA chelation of Mg to block ATP hydrolysis (bar 8 vs 16) or by employing hyrolysis-resistant ATPγS instead of ATP (bar 8 vs 9) and is less pronounced at low ATP levels (Figure 6C, bars 2 or 4 vs 8),).

**Figure 6.**
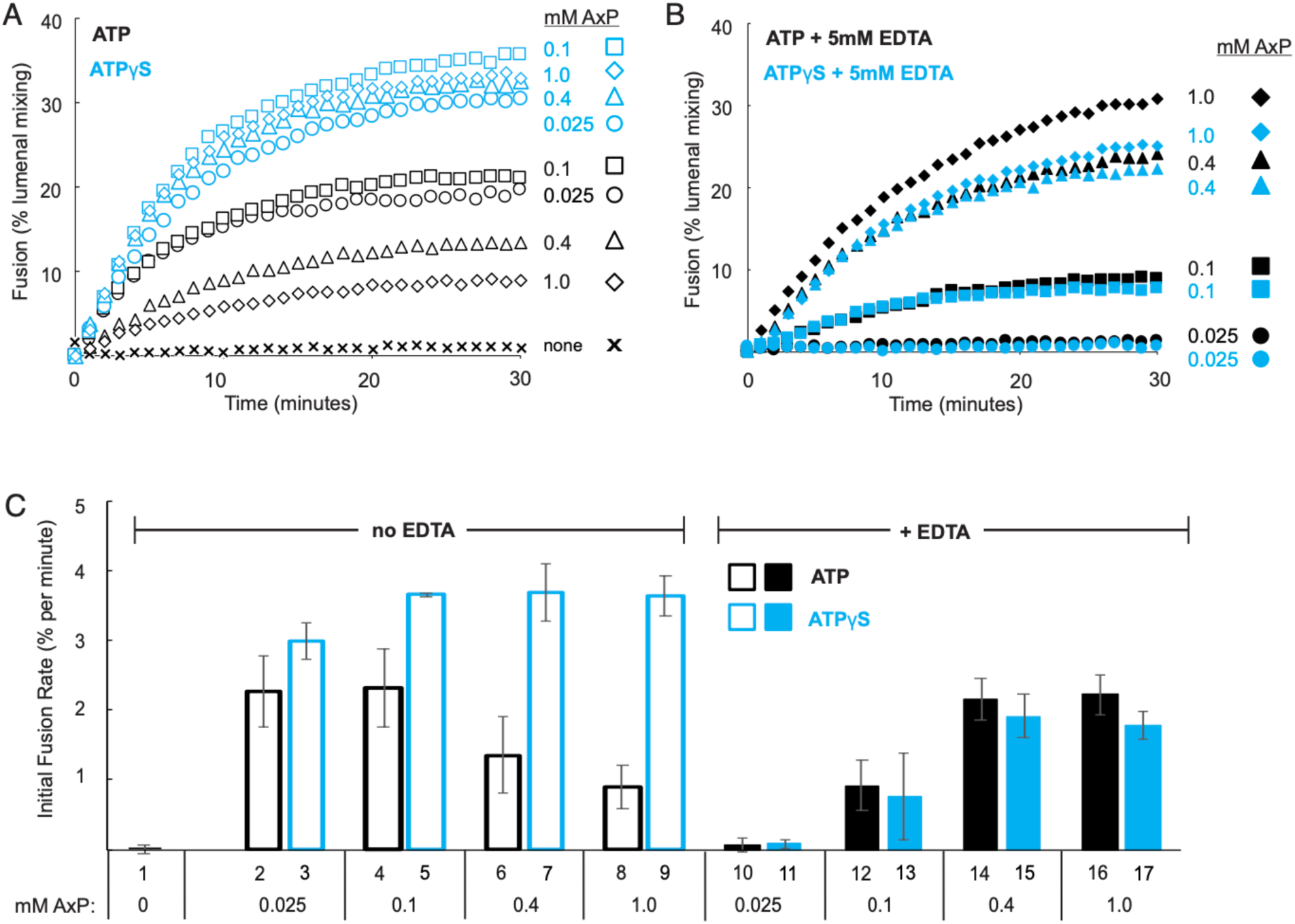
ATP and ATPγS are equipotent as Sec18 ligands for Sec17/Sec18-mediated fusion, but hydrolysis of ATP inhibits fusion. Fusion incubations were performed with 400 nM Qc3Δ, 50 nM HOPS, 0.1 mM Sec17, and either no adenine nucleotide (x) or the indicated levels of MgATP (black symbols) or MgATPγS (blue). Proteoliposomes (8 μl) were prewarmed in separate wells from a mixture of all soluble components (9 μl). After 10 minutes at 27°C, 3 μl of either Rb150 buffer or of 40 mM EDTA in Rb150 was added to the soluble components by a multichannel pipettor, mixed, and 8 μl of the 12 μl mixture was transferred to the well with proteoliposomes, mixed with the tips on the multichannel pipettor, and fusion recording begun. (A) No EDTA, (B) 5 mM EDTA, (C) Median and standard deviations of the initial fusion rates from triplicate assays.

### Sec17 release without SNARE disassembly

Sec18, Sec17 and ATP disassemble SNARE complexes (Söllner et al., 1993). Is this the basis for the inhibition of Sec17/Sec18-mediated fusion by ATP hydrolysis? Fusion incubations were performed with Rab/R and Rab/Qab proteoliposomes with HOPS, Qc3Δ, Sec18, 0.1mM Sec17, and either ATP or ATPγS. Much less fusion was seen with ATP than with ATPγS (Figure 7A, filled vs open circles). Membranes from these incubations and from control incubations which either lacked HOPS or lacked the R-proteoliposomes were solubilized in modified RIPA buffer (Orr and Wickner, 2023) and mixed with bead-bound antibody to Qa to allow analysis of Qa-bound proteins (Figure 7B). There was no difference between incubations with ATP or ATPγS in the Qa-bound R, a measure of trans-SNARE complex, or Qa-bound Qb or Qc, and thus no ATP-dependent disassembly of SNARE complex (Figure 7B, lanes 2, 3). However, there was less Qa-bound Sec17 from incubations with ATP (lane 3). Control incubations showed that the binding of R to Qa relied on HOPS (lane 1) and that the binding of Sec17 to Qa (lane 2) relied on both HOPS (lane 1) and on the Rab/R proteoliposomes (lane 4), and thus reflected Sec17 associated with *trans*-SNARE complex. With limiting Sec17 (0.1 mM), ATP supports selective Sec17 release from trans-SNARE complexes with accompanying loss of fusion.

**Figure 7.**
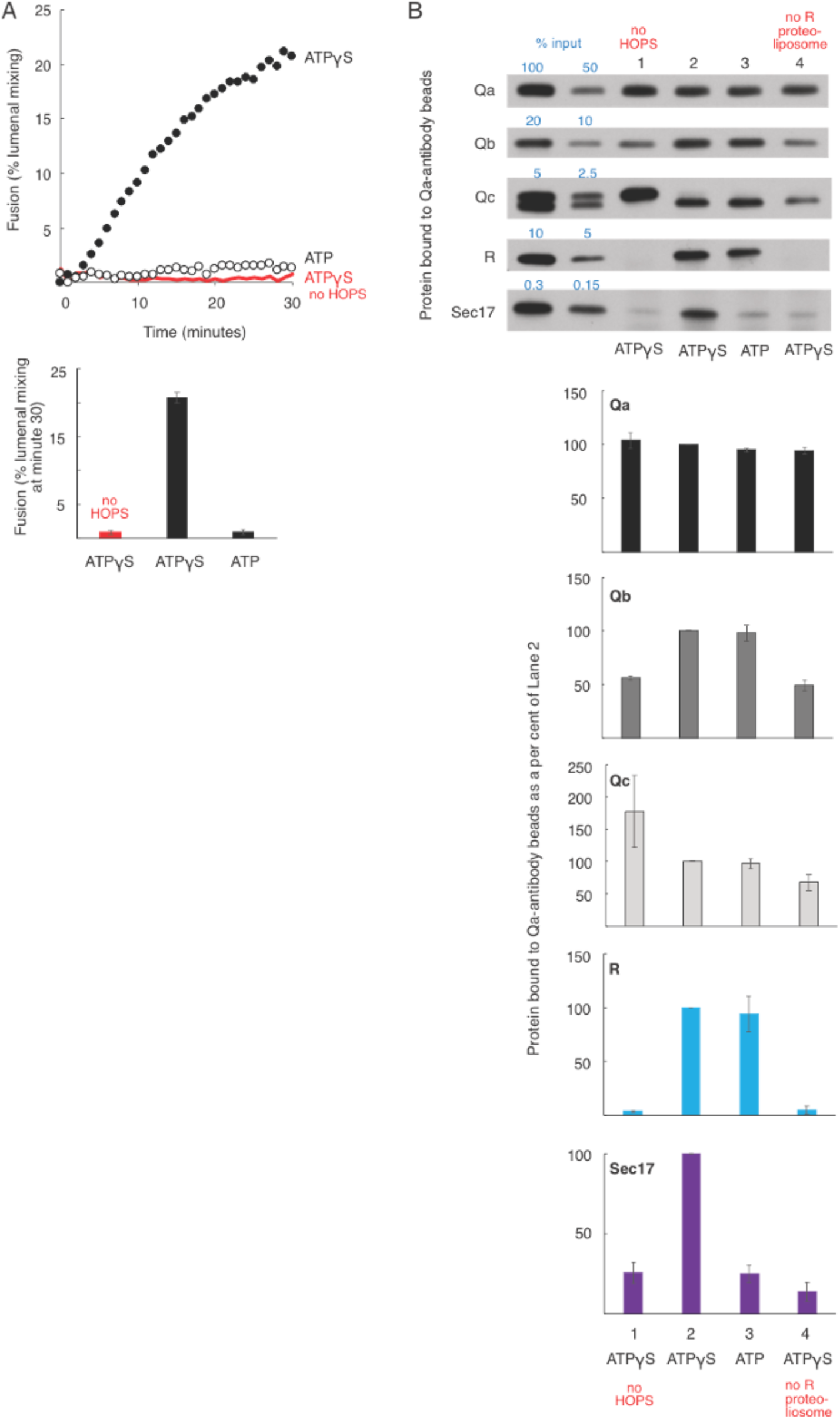
At limiting Sec17, Sec18 and ATP displace Sec17 from the *trans*-SNARE complex without disassembling that complex. Fusion reactions were conducted as described in Materials and Methods in 20 µl volumes with Rab/R + Rab/Qab proteoliposomes, 0.4µM Qc3Δ, 50 nM HOPS, 0.1µM Sec17, and 0.4µM Sec18 with 1 mM Mg:ATP or 1 mM MgATPyS. In addition to the 20 µl reactions in the 384 well plate, identical larger scale reactions were conducted simultaneously in PCR strips. After reading FRET for 30 minutes in the plate reader, 72 µl of each reaction were added to 1.5 ml tubes containing 15 µl Protein A magnetic beads, 7.5 µg affinity purified antibody to Qa, and 5 µM GST-His-Nyv1 as a competitor, all in RIPA Buffer in a total volume of 225 µl. Reactions were nutated end over end for 2 hours at 4°C, then washed 3 times in RIPA Buffer. Bound proteins were eluted by boiling for 2 minutes in sample buffer with B-mercaptoethanol and analyzed by SDS page gel followed by immunoblotting. The results of triplicate assays were quantified with UN-SCAN-IT software (Orem, UT) and are shown here with error bars. (A) With defective SNARE zippering, Sec17/Sec18-driven fusion relies on HOPS and non-hydrolysable ATP. (B) Trans-SNARE complex remains assembled with either ATP or ATPγS, while Sec17 is absent from the complex in the presence of hydrolysable Sec18:Mg:ATP.

### With all wild-type SNAREs and unimpeded zippering, lipidic impairment of fusion is bypassed by Sec17, Sec18, and adenine nucleotides

To test whether stimulation by Sec17 and Sec18 is a general feature of impaired fusion or is only seen with mutant SNAREs, proteoliposomes were prepared as before with vacuolar mixed lipid headgroup composition and with the same distributions of both Rab and R– and Q-SNAREs but with palmitoyl, oleoyl (PO; 16:0, 18:1) fatty acyl chains instead of the more fluid 18:2, 18:2 fatty acyl chains. PO fatty acyl chains impair reconstituted vacuolar fusion (Zick and Wickner, 2016). The slow fusion of proteoliposomes with PO fatty acyl chains (Figure 8A, open black squares) was hardly affected by 0.1 or 1.0 μM Sec17 alone (open black diamonds or circles) but was stimulated by Sec18 without Sec17 with either ATP or ATPγS (red or blue filled squares). There was no further stimulation by higher concentrations of Sec18 alone (Figure 8B). Optimal fusion required Sec17 as well as Sec18 (Figure 8A, filled diamonds and triangles; Figure 8B, open black squares) but needed no more than 10 nM Sec17 (Figure 8A, filled blue and red triangles and diamonds). These findings contrast with those seen with defective zippering, where Sec17 alone (but not Sec18) gave some fusion support (Figure 3A) and where high Sec17 concentrations were needed for optimal fusion with Sec18 and ATP (Figure 3B).

**Figure 8.**
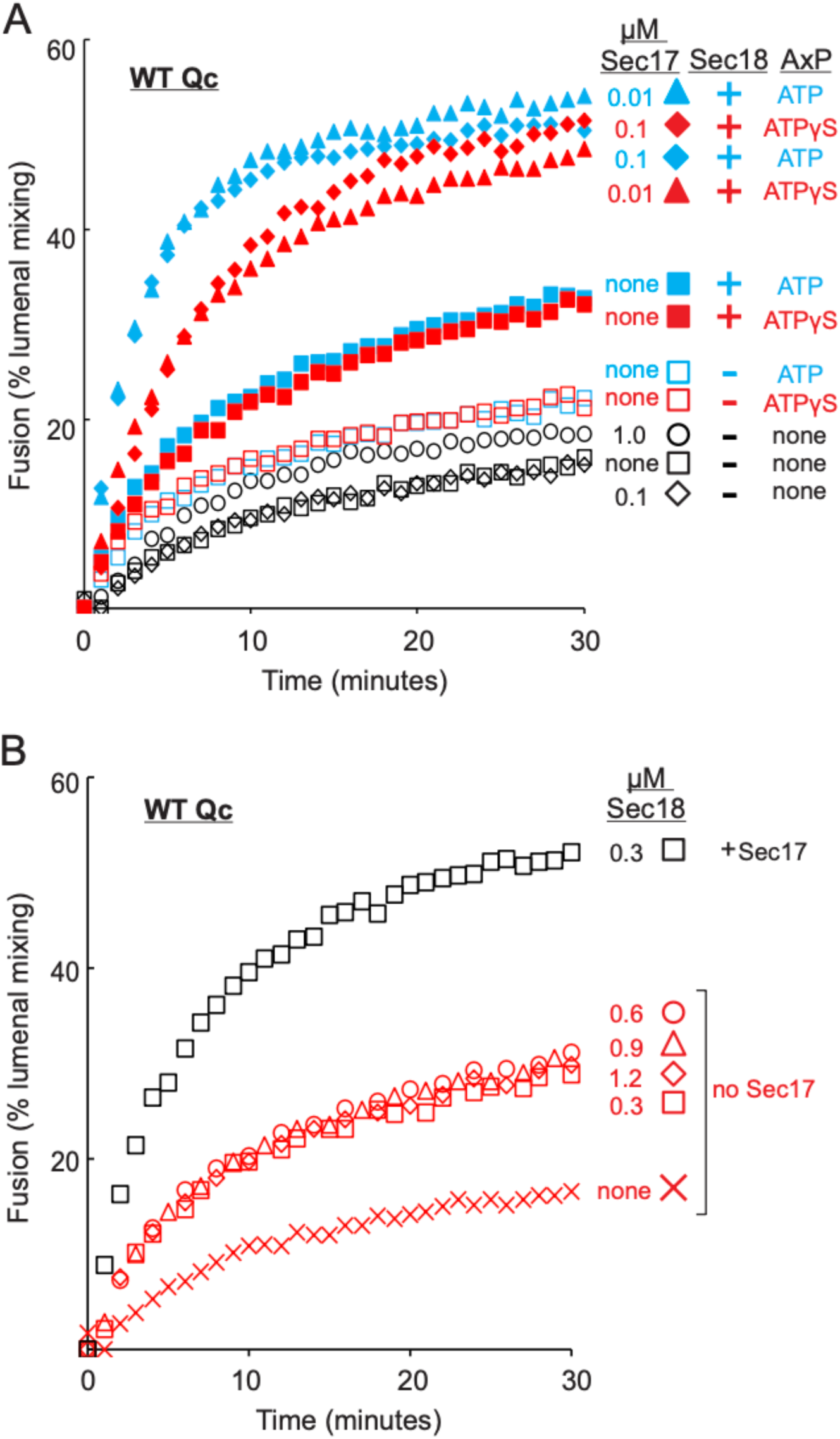
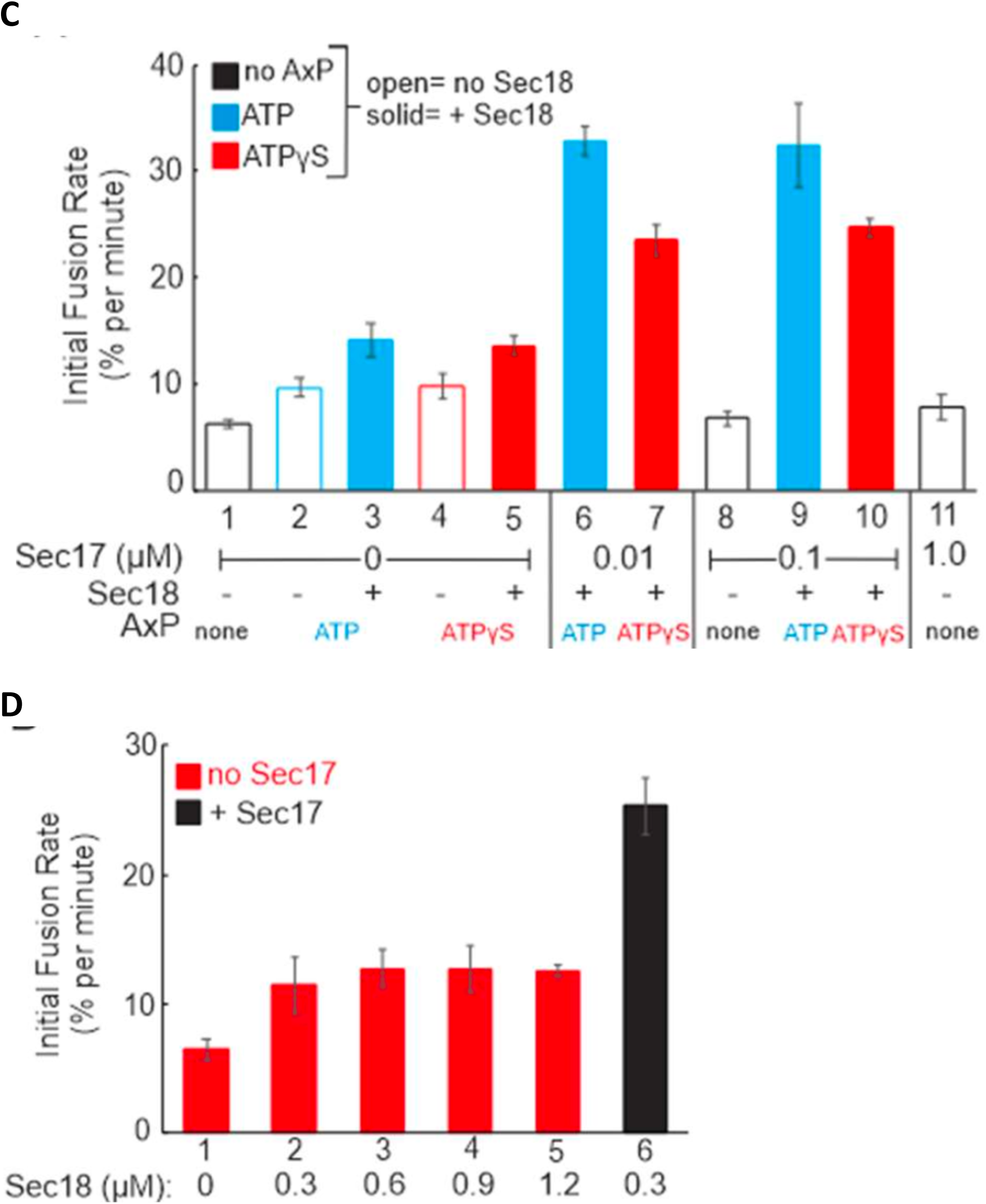
Fusion with all wild-type SNAREs but limited by the fatty acyl composition of the bilayer is enhanced by Sec18 and either ATP or ATPγS, with further enhancement by Sec17. Rab/R and Rab/QaQb proteoliposomes were prepared as described in Methods, but with 16:0, 18:1 (palmitoyl, oleoyl or PO) PC, PE, PA, and PS, a 1:4,000 molar ratio of Ypt7-tm to lipids, a 1:10,000 molar ratio of R to lipids, and a 1:2,000 molar ratio of Qa and Qb to lipids, as per Zick et al. (2015). (A) Fusion incubations had 50 nM HOPS, 200 nM Qc, and 0.01, 0.1, or 1.0 μM Sec17, 0.3 μM Sec18, and 1 mM MgATP or MgATPγS as indicated. (B) Fusion incubations as above, with the indicated levels of Sec17 and Sec18. (C, D) Mean fusion rates and standard deviations for triplicate repeats of panels A, B.

## Discussion

Sec17 is present at 12,000 copies per cell (Ho et al., 2018), and the cytoplasmic volume per cell is 20 μm^3^ (Uchida et al., 2011). The Sec17 cytoplasmic concentration is therefore approximately 1 μM, though the free concentration may be less as some Sec17 is likely associated with SNAREs or other proteins. Whether with fluid membranes and wild-type SNAREs (Figure 2), with SNAREs impaired for zippering-derived energy for fusion (Figures 3-7), or with wild-type SNAREs but stiff fatty acyl chains (Figure 8), Sec17 and Sec18 are important for fusion. High Sec17 levels can function without Sec18, but Sec18 stimulation allows lower Sec17 concentrations to function effectively for fusion (Figures 3, 8). With impaired zippering and at limiting Sec17 levels, Sec18-mediated ATP hydrolysis blocks fusion by releasing Sec17 from *trans*-SNARE complexes without concomitant SNARE disassembly (Figure 7). With wild-type SNAREs and unimpeded SNARE zippering, fusion can also be suppressed by the fatty acyl chain composition of the major lipids (Figure 8). This suppression is relieved by the synergistic action of Sec17 and Sec18, but with several clear distinctions from Sec17/Sec18 function when deletion of the C-terminal region of one or more Q-SNAREs diminishes the energy from zippering (Song et al., 2021). Sec17/Sec18-mediated fusion with Q-SNARE C-terminal truncation can be relieved by high concentrations of Sec17 alone (Schwartz and Merz, 2009; Zick et al., 2015; Schwartz et al., 2017), but not by Sec18 alone. Sec17 alone at concentrations up to 1 μM has little effect on fusion with PO fatty acyl chains (Figure 8A), but Sec18 alone stimulates this fusion (Figure 8A, B). With 0.1 μM Sec17, Sec17/Sec18-mediated fusion with a truncated Qc SNARE is blocked by ATP hydrolysis-driven selective Sec17 release (Figure 7). With wild-type SNAREs and PO lipids, Sec18-dependent fusion is equivalent with ATP or its hydrolysis-resistant analog ATPγS (Figure 8A).

We present a speculative model of how Sec17, Sec18, and ATP may support rapid fusion in these two conditions where sufficient fusion energy is not available from SNARE zippering alone. The 20s structure of SNAREs, NSF/Sec18, and SNAP/Sec17 (Zhao et al., 2015) shows that several side by side Sec17 molecules can form a cylinder wrapped around the 4-SNARE complex, as illustrated for *trans*-SNARE complexes in Figure 1. Each Sec17 molecule is bound to Sec18, lipid, SNAREs, and the other Sec17 molecules in the 20s structure. Each Sec17 is engaged through its two C-terminal leucines with Sec18 at the membrane-distal C-terminus of Sec17, with the membrane bilayer through the Sec17 N-domain apolar loop, and with SNAREs through basic residues in the Sec17 center (Figure 1, cross section view). Each Sec17 has an acidic edge and a basic edge. In the complete 20s structure, both edges of each Sec17 are electrostatically bound to its neighbor. With incomplete SNARE zippering and at low and limiting Sec17 concentrations, *trans*-SNARE complexes will not have a full complement of bound Sec17. Some Sec17 molecules would be weakly bound if they were not associated with a Sec17 neighbor at all or with only one neighboring Sec17. ATP hydrolysis by Sec18 may displace these Sec17. With fewer Sec17 and with each lacking contact with the unzippered C-terminal region of the SNARE domains, there may be insufficient force applied to SNAREs during Sec17 displacement to disassemble the 4-SNARE complex. Further studies are needed to test this model.

Sec17 and Sec18 stimulate fusion with fluid membranes and wild-type SNAREs (Figure 2) but are more strictly required under a variety of conditions which otherwise limit fusion. These include C-terminal truncation of any of the Q-SNAREs to arrest zippering (Schwartz and Merz, 2009; Song et al., 2021), swap of the juxtamembrane domains of the R and Qa SNAREs (Orr et al., 2022), anchoring of the Qb SNARE instead of the Qa SNARE (Wickner et al., 2023), and stiff and less fluid lipids with 16:0, 18:1 fatty acyl chains (Figure 8, and Zick et al., 2015). Sec17 without Sec18 can stimulate the fusion of proteoliposomes with R and Qa that have swapped juxtamembrane regions (Orr et al., 2022) and Sec18 without Sec17 stimulates fusion with wild-type SNAREs at low HOPS (Orr and Wickner, 2024). It is unclear how Sec18 can stimulate fusion without Sec17, especially since that stimulation does not need energy from ATP hydrolysis (Figure 8A). Sec18 can interact directly with SNAREs (Zick et al., 2015), but is only known to bind them at a substantial distance from the membrane (Zhao et al., 2015). Sec18 is large, and bulk itself might contribute to membrane bending (Agostino et al., 2017).

Our current working model of vacuole fusion begins with vacuoles whose SNAREs had been disassembled by Sec17, Sec18 and ATP after a prior round of fusion. Vacuoles are tethered by Vps39 and Vps41 subunits of HOPS binding to the Rab Ypt7 on each fusion partner membrane (Stroupe et al., 2006; Hickey and Wickner, 2010; Brett et al., 2008; Bröcker et al., 2012). When Rab-bound HOPS also engages both phosphatidylinositol-3-phosphate and R-SNARE in *cis*, it is activated to catalyze the assembly of SNARE complex in trans between the bound R and the Q-SNAREs from the apposed, tethered vacuole (Torng and Wickner, 2020; Song et al., 2021). As SNARE zippering proceeds, the R and Qa SNARE domain residues that are in a groove on the surface of the HOPS subunit Vps33 (Baker et al., 2015) turn towards each other in the center of the 4-helical bundle. This may weaken the affinity of HOPS for the SNAREs. Several Sec17s form a transient complex with Sec18, and this complex binds directly to membranes by the product of the affinity of each Sec17 apolar loop for lipid (Orr and Wickner, 2022). Sec17:Sec18 then assemble with the partially-zippered SNAREs to form what might be termed a “*trans*-20s complex” (Figure 1). Sec17 may promote HOPS release (Collins et al., 2005; Schwartz et al., 2017) and promote zippering (Ma et al., 2016; Song et al., 2021), but its membrane-inserted N-domain apolar loops also directly promote the lipid rearrangements of fusion (Song et al., 2021). Sec18 also directly promotes fusion in both Sec17-dependent and Sec17-independent manners (Figure 2, 3, and 8), though these are not understood at a molecular level. The purpose of this model is to suggest critical experiments. Many aspects of fusion remain elusive; for example, it is unclear why the Qb membrane anchor is dispensible for fusion while the Qa anchor is essential (Wickner et al., 2023), why swap of the juxtamembrane regions between R and Qa SNAREs blocks zippering-driven fusion (Orr et al., 2022), or which combination of the Sec18 affinities for SNAREs (Zick et al., 2015), Sec17 (Weidman et al., 1989), and HOPS (Orr and Wickner, 2024) allow it to support fusion from such a distance from the lipid bilayer (Figure 1).

## Materials and Methods

### Materials and Proteins

Lipids were purchased from Avanti Polar Lipids (Alabaster, AL), Echelon Biosciences (Salt Lake City, UT), and Sigma-Aldrich (St. Louis, MO). β-octylglucoside was from Anatrace (Maumee, OH). Biotinylated phycoerythrin and Cy5-streptavidin were purchased from Life Technologies Corp (Eugene, OR) and SeraCare (Milford, MA), respectively. GTP, ATPγS, and Histodenz were from Sigma-Aldrich. Biobeads SM2 were from Biorad (Hercules, CA). The vacuolar SNAREs (Mima et al., 2008), Rab Ypt7-tm with a C-terminal transmembrane anchor instead of a prenyl anchor (Song et al., 2020), Sec17 (Schwartz and Merz, 2009), Sec18 (Haas and Wickner, 1998), HOPS (Zick and Wickner, 2013), Sec17-FSMS (Zick et al., 2015), Qc3Δ (Schwartz and Merz, 2009), and GST-His_6_-Nyv1 (Izawa et. al, 2012), used as a competitor in immunoprecipitation assays of Figure 7, were purified as reported, frozen in small aliquots in liquid nitrogen, and stored at –80°C.

### Proteoliposome preparation

Proteoliposomes were prepared as described (Orr and Wickner, 2022). Briefly, β-octylglucoside in methanol was mixed with chloroform solutions of lipids of vacuolar-mimic composition (18:2, 18:2 PC, PE, PS, and PA; soy PI, ergosterol, diacylglycerol, and synthetic diC16-PI3P mole % of 46, 18, 18, 4.4, 2, 8, 1, and 1% respectively), dried under a stream of nitrogen followed by 3 h speedvac, rehydrated to 10 mM lipid in 2.5x concentrated Rb150 [Rb, reaction buffer, is 20 mM HEPES/NaOH, pH 7.4, 150 mM NaCl, 10% glycerol] with 2.5 mM MgCl_2_. Portions of 400 μl were transferred to 2 ml screw-cap glass vials, topped with argon, and stored at –80°C. Prior to use, each was thawed and nutated for at least 30 min at room temperature, then placed on ice. SNAREs and the Rab Ypt7-tm were added to 1:32,000 and 1:8000 molar ratio to lipids, respectively, then 0.25 ml of either 32 µM Cy5-streptavidin or 16 µM biotinylated phycoerythrin, and water to 1 ml. This 1 ml mixed micellar solution was placed in 25 kDa cutoff dialysis tubing, sealed, and dialyzed with stirring for at least 16 h at 4°C in the dark against 200 ml Rb150 + 1 mM MgCl_2_ and 1 g Biobeads SM2. Dialysate was mixed with 1 ml of 70% Histodenz in iso-osmolar Rb150 + 1 mM MgCl_2_ in an ultraclear SW60 tube (Beckman-Coulter, Brea, CA), then overlaid with 1.6 ml 25% Histodenz in Rb150 + 1 mM MgCl_2_ and finally 800 μl Rb150 +1 mM MgCl_2_. After centrifugation for 90 min at 3°C, 55,000 rpm, floated proteoliposomes were recovered from the 0/25% interface. Proteoliposomes were assayed for lipid phosphorus, the diluted with Rb150 + MgCl_2_ to 2 mM. Aliquots of 30 μl in 250 μl tubes were frozen by immersion in liquid nitrogen and stored in liquid nitrogen.

### Assay of fusion

Proteoliposome fusion was assayed as described in detail (Orr and Wickner, 2023). In brief, Rab/R and Rab/QaQb proteoliposomes (0.44 mM lipid each), nonfluorescent streptavidin (10 μM), EDTA (1 mM), and GTP (50 μM) were mixed on ice, incubated for 10 min at 27°C, then returned to ice and mixed with 3 mM MgCl_2_ to complete the loading of GTP onto the Rab. A separate mixture was prepared with each of the soluble fusion reaction components, such as the Mg salt of an adenine nucleotide, Qc or Qc3Δ, HOPS, Sec17, and Sec18 as noted in each figure. After 10 min separate preincubation of 8 μl of the proteoliposomes and 11 μl of the mixed soluble components, a multichannel pipettor was used to deliver 8 μl of mixed soluble components and to mix them with the proteoliposomes. FRET between the Cy5 and phycoerythrin fluorophores was assayed as a measure of fusion at 1 min intervals with a Molecular Devices SpectraMax Gemini plate reader, Ex 565 nm, Em 670 nm, cutoff 630 nm.

## Acknowledgements

This work was supported by grant 2R35GM118037 from the National Institute of General Medical Sciences. We thank Axel Brunger and Gus Lienhard for fruitful discussions and suggestions.

